# Multi-scale classification decodes the complexity of the human E3 ligome

**DOI:** 10.1101/2025.03.09.642240

**Authors:** Arghya Dutta, Alberto Cristiani, Siddhanta V. Nikte, Jonathan Eisert, Ramachandra M. Bhaskara

## Abstract

E3 ubiquitin ligases are key regulators of protein homeostasis, targeting specific proteins for degradation via the ubiquitin-proteasome system (UPS). They provide crucial substrate specificity, making them promising candidates for the design of novel therapeutics. This work presents a comprehensive, annotated dataset of high-confidence catalytic human E3 ligases, termed the “E3 ligome”. Integrating disparate data from various granularity layers, including protein sequence, domain architecture, 3D structure, function, localization, and expression, we learn an emergent distance metric, capturing authentic relationships within this heterogeneous group. A weakly-supervised hierarchical classification framework identifies conserved features of E3 families and subfamilies, consistent with RING, HECT, and RBR classes. This classification explains functional segregation, identifies multi-subunit and standalone enzymes, and integrates substrate and small molecule interaction networks. Our analysis provides a global view of E3 biology, opening new strategies for drugging E3-substrate networks, including drug re-purposing and designing new E3 handles.

## Introduction

Cells constantly modulate their proteomes in response to physiological and environmental changes. The timely removal and turnover of cellular proteins is integral to protein homeostasis (*1*). In eukaryotes, individual proteins, complexes, and large assemblies are degraded via either autophagy or the ubiquitin-proteasome system (UPS) (*2*). In mammalian cells, approximately 80% of the cellular proteome is degraded through the UPS (*1*). In this pathway, the designated protein cargo is tagged with ubiquitin (Ub) molecules through a series of enzymatic reactions, marking them for degradation by the proteasome (*3*). Following the action of E1 and E2 enzymes, the E3 ligase brings both the E2–ubiquitin complex and the substrate protein in proximity, allowing the transfer of Ub from the E2 enzyme to a lysine residue on the target protein (*4, 5*). This process is often repeated (polyubiquitination), resulting in substrates with distinct types of Ub-chains. In UPS, for instance, K48-linked Ub-chains are recognized by Ub-binding domains (UBDs) on 19S proteasomal particles, initiating the degradation of substrates (*1*). In autophagy, ubiquitination often serves as a necessary condition for identifying substrates, conferring specificity (*6*). Cargo components, damaged organelles, and intracellular pathogens targeted for degradation are often ubiquitinated. Further, autophagy receptors are enriched in UBDs to recognize modified cargo components (*7*) or themselves strongly ubiquitinated to trigger aggregation of protein assemblies in the cytosol and organellar membranes (*8, 9*), thus enhancing autophagic flux.

E3 ubiquitin ligases confer substrate specificity for ubiquitination. They recognize distinct targets, operate in diverse cellular locations, and exert spatial control of protein turnover (*10, 11*). In addition to controlling homeostatic processes, E3 ligases regulate immunity and inflammation pathways (*12, 13*). Given their tissue-specific expressions and association with developmental and metabolic syndromes, including cancer progression, E3 ligases have emerged as promising candidates, particularly for drugging previously undruggable targets (*14*). In stark contrast to E1 (∼10) and E2 enzymes (∼50), a substantial number of E3 ligases (∼600) have been recognized in humans (*15,16*). This count of putative E3s stems from various investigations: Li et al. (*17*) identified 617 potential human E3-encoding genes by conducting a genome-wide search to detect RING (Really Interesting New Gene) finger catalytic domains using hidden Markov models. Subsequently, Deshaies and Joazeiro (*18*) characterized ∼300 RING and U-box E3 ligases, while Medvar et al. (*19*) documented ∼377 E3 ligases, with a primary focus on confirmed catalytic activity. Despite these efforts, many human E3 ligases have been only partially characterized. A significant fraction remains unexplored and hypothetical or unknown (*20*). To date, those studied exhibit extensive heterogeneity in their sequence, domain composition, 3D structure, subcellular localization, and tissue expression, establishing them as one of the most diverse classes of enzymes. Furthermore, several E3 ligases function as multi-subunit complexes with varied substrate specificities modulated by specific receptors, adaptors, and scaffold proteins (*21*). The extensive variety and large numbers of E3 ubiquitin ligases create a bottleneck for pattern recognition and large-scale study. Therefore, detailed characterization and analysis of the human E3 ligome—the complete set of E3 ubiquitin ligases encoded by the human genome—is essential for a comprehensive understanding.

The current classification of the E3 ligases—based on the ubiquitin-transfer mechanism— categorizes them into three main classes: RING (Really Interesting New Gene), HECT (Homologous to the E6AP Carboxyl Terminus), and RBR (RING-Between-RING) classes (*15*). This classification drastically oversimplifies the mechanistic diversity of E3 ligases, compels the grouping of enzymes with hybrid characteristics, and fails to accommodate emerging information on new and atypical ligases, limiting its overall utility (*18*). A multi-scale classification of the human E3 ligome offers a unique solution to tackle the complexity and remarkable diversity inherent in these enzymes at various scales. This organized approach can provide more accurate and functional groupings crucial for a nuanced understanding of different E3 ligase families. Further, novel patterns detected help trace evolutionary relationships more effectively, revealing conserved elements and adaptive changes that are not evident. Furthermore, mapping essential information such as functional diversity, substrate-specificities, and druggability onto the classification provides a global view, guiding specific and directed investigations to fill in the missing information.

Here, we systematically catalog all E3 ubiquitin ligases to build a comprehensive and manually curated human E3 ligome. We then encode the relationships between high-confidence E3 ligases using multiple distance measures at various granular layers spanning the molecular- and the systems-level organization. By amalgamating selected distance measures from multiple layers into an optimized emergent distance metric, we group all human E3 ligases into distinct families and subfamilies. Our classification delineates features and patterns specific to E3 ligase families, providing insights into their organization. We demonstrate the utility of this unbiased classification by mapping the existing state of knowledge on E3 ligase domain architecture, 3D structure, function, substrate networks, and small molecule interactions to gain generic and family-specific insights. The multiscale classification framework developed here offers a comprehensive roadmap to navigate the vast landscape of E3 ligase biology, laying the groundwork for future therapeutic applications.

## Results

### Assembly of the human E3 ligome

To comprehensively identify all E3 ligases in the human genome, we conducted a census using datasets from previously published studies and public repositories. By visualizing their overlaps, we found that all existing datasets were largely inconsistent (**Fig. 1a** and **Fig. S1a**). Most strikingly, only 99 proteins were consistently categorized as human E3 ligases from all eight datasets. The low overlap in these datasets reflects the diverse approaches and often variable and fuzzy definitions used to collate E3 systems (**Table S1**). We resolved these conflicts by clearly defining the catalytic components of E3 systems, i.e., polypeptide sequences containing one or more catalytic domains (*C* = {*d*_*c*_}, see methods). Using this objective criterion (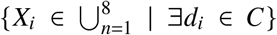**Table S2**) facilitated proper annotation and targeted analysis of E3s. We found that 462 polypeptide sequences, across all datasets 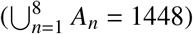, contain at least one catalytic domain constituting the curated E3 ligome (**Fig. 1b** and **Fig. S1b**).

**Figure 1:**
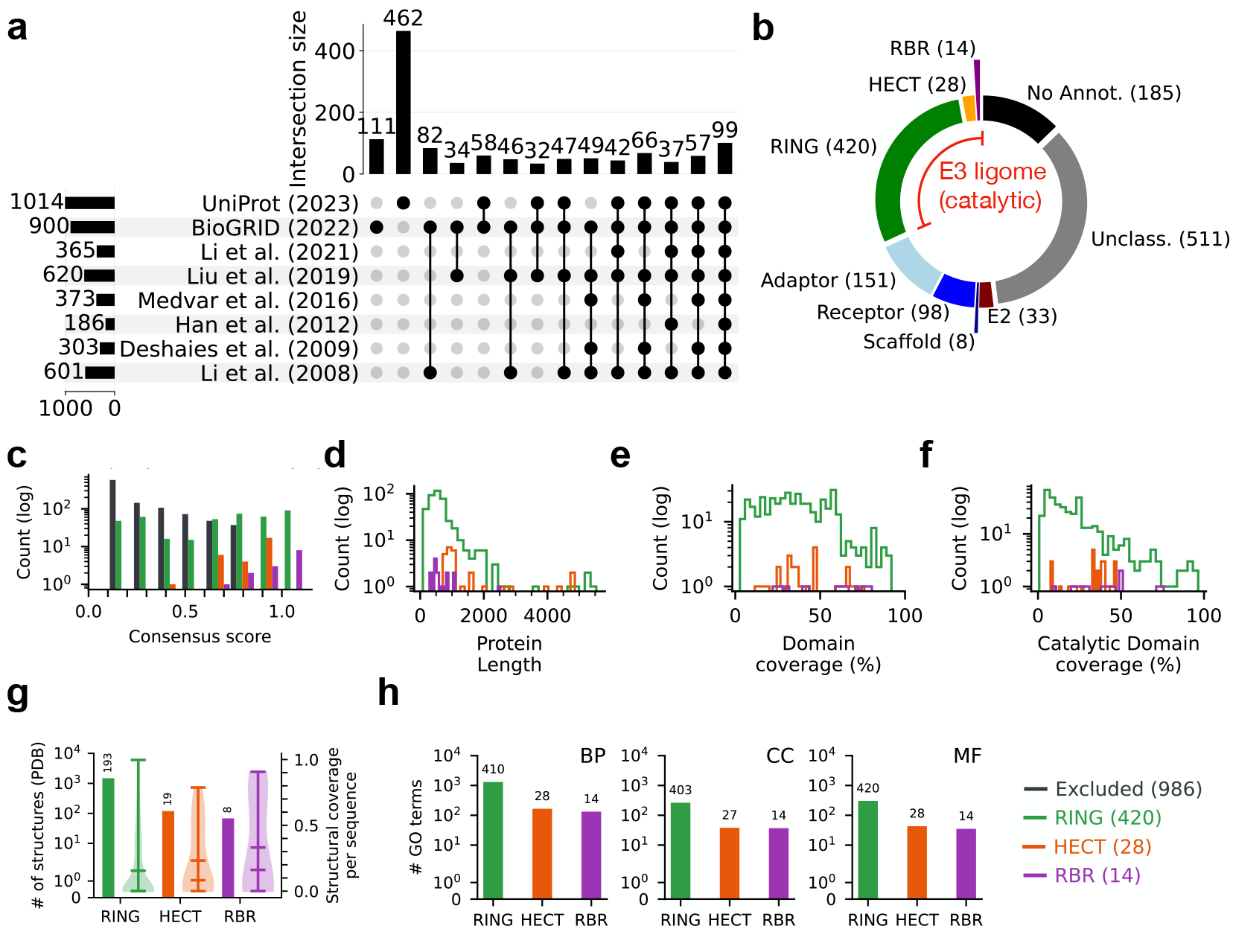
Diversity of the human E3 ligome. **(a)** A visualization showing the intersections of eight E3 ligases datasets (*A*_1_, ···, *A*_8_) obtained from existing literature and public repositories. The matrix layout for all intersections of individual datasets is sorted by size. Filled circles and their corresponding bars indicate sets that are part of the intersection and their sizes, respectively. Individual proteins (*X*_*i*_) from the all eight datasets 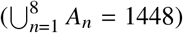 annotated with one or more domains, *d*_*i*_, belonging to a set of well-studied catalytic compone nts of E3 enzymes (*C* = {*d*_*c*_}) were compiled to form the high-confidence E3 ligome,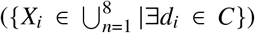. **(b)** Pie chart showing the extent of protein annotations and filtering to identify the catalytic components of the human E3 ligome. **(c)** The histogram of consensus scores for each entry quantifies their distribution among RING (420), HECT (28), and RBR (14) classes. The distribution of **(d)** protein lengths and annotation coverage for **(e)** all domains and **(f)** catalytic domains highlights the heterogeneity of the E3 ligome. **(g)** Distribution of structural coverage of the E3 ligome at class-level. **(h)** The total number of unique GO terms associated with E3 classes indicates their functional vista under biological process (BP), cellular component (CC), and molecular function (MF) ontologies.

To substantiate our curation process, we defined a consensus score for each protein based on its presence in various source datasets (**Fig. 1c**). We found that the HECT and RBR classes of E3 ligases showed high agreement across datasets (confidence score ≥ 0.6; orange and purple bars). The RING class (green bars) had a broad distribution of consensus scores indicative of annotation challenges. However, the most significant discrepancy among the datasets (confidence score ≤ 0.25) was due to misannotated proteins. E1, E2, and other non-catalytic components of E3 systems, such as receptors, scaffolds, and adaptor proteins, were often merged with E3 ligases (**Fig. 1b**). Furthermore, several proteins obtained from UniProt and BioGRID using keyword-based searches (**Fig. S1c**) have low consensus scores and remain unclassified and unannotated, excluding 986 proteins from the curated E3 ligome (**Fig. 1c**, black bars). Our approach thus minimized false positives and provided high-confidence catalytically active E3s.

To get an initial assessment and quantify the diversity of the human E3 ligome, we mapped the sequence, structure, and functional features of individual E3s corresponding to well-known E3 classes (RING, HECT, and RBR). We found that the length distribution of the E3s is broad, ranging from 100 to 5000 residues (mean size = 635 residues; **Fig. 1d**). The average fractional coverage of E3s annotated with unique domains is 37%, 42%, and 53% for RING, HECT, and RBR classes, respectively (**Fig. 1e**). Furthermore, on average, the RING, HECT, and RBR domains span 23%, 31%, and 39% of their total lengths, respectively (**Fig. 1f**). By mapping information from the Protein Data Bank (PDB), we found 1675 distinct structures representing RING, HECT, and RBR– containing proteins (1488+119+68), providing partial structural information for 47% (193+19+8) of the E3 ligome (**Fig. 1g**). Analysis of AlphaFold models revealed that for most E3s, the coverage of structured domains is high, and the amount of intrinsic disorder is generally low (pLDDT ≤ 50 covering only ≤ 10% E3 length; **Fig. S1d**). We quantified the functional diversity of the E3 ligome by retrieving the unique Gene Ontology (GO) annotations corresponding to Biological Processes (BP), Cellular Component (CC), and Molecular Function (MF). We annotated 96–100% of the E3s with unique GO terms (**Fig. 1h**). The number of distinct GO terms captured the diversity of functional assignments attributed to the three E3 classes.

### Metric learning for classification of the human E3 ligome

To study the organization and relationships of proteins within the human E3 ligome, we attempted to classify these enzymes using multiple sequence alignment (MSA) followed by phylogenetic tree construction. However, we obtained a low-quality MSA with numerous gaps (**Fig. S2a**), primarily due to (i) high sequence divergence, (ii) numerous proteins with uneven length distributions, (iii) inadequate alignment of conserved, catalytic domains, and (iv) an extensive repertoire of domain architectures (**Fig. S2b**).

To capture the complex relationships within the human E3 ligome, we used a machine-learning approach to learn an emergent distance measure. Using a linear sum model, we combined multiple distance measures with optimal weights to reproduce class-level organization (partial ground truth) in hierarchical clustering (**Fig. 2a**). We first computed twelve pairwise distance matrices for all E3 ligases (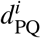 where *i* = {1, ···, 12}, for all E3s P and Q ∈ E3 ligome; 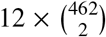 distances) across distinct granular layers: primary sequence, domain architecture, 3D structure, function, subcellular localization and expressions (see methods). These distances between ligase pairs are widely distributed and capture their relationships across distinct molecular- and systems-level hierarchies (**Fig. 2b**). Interestingly, most distance measurements showed low correlations (**Fig. 2c**), suggesting that they capture largely orthogonal information from the distinct granularity layers. Only the three domain architecture-based distances which quantify domain composition 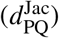, domain order 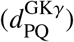, and domain duplication 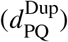 are highly correlated (Pearson *r* ≥ 0.5).

**Figure 2:**
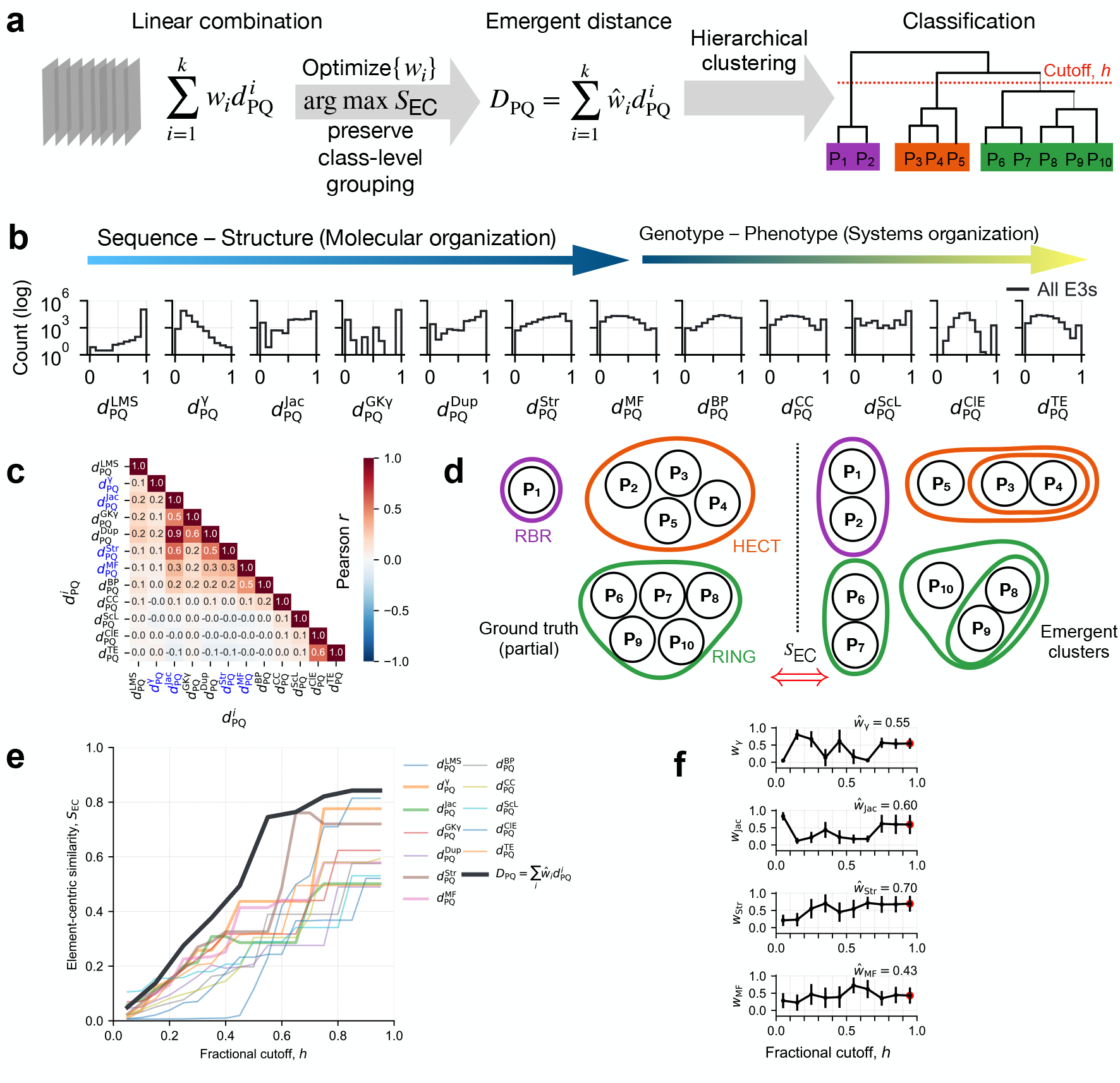
Metric learning for E3 ligases. **(a)** Schematic of the metric learning process. **(b)** Distribution of various pairwise distance measures spanning the molecular and systems level organization. **(c)** Pearson correlation of distance measures indicate orthogonality, mostly *r* ∈ (*−*0.3, 0.3). Distances based on sequence alignment, domain composition, 3D structure (catalytic), and molecular function (marked in blue) are combined into an emergent distance (*D*_PQ_) with appropriate weights. **(d)** By maximizing element-centric similarity, a measure of the overlap of emergent hierarchical clusters (right) with the ground truth (left) **(e)** evaluates individual metrics and their linear combinations. **(f)** Regression weights (mean ± S.D.) corresponding to the four relevant distances as a function of fractional tree cutoff *h*. 100 clusters with largest *S*_EC_ were sampled at each value of *h* to estimate the mean and S.D.

Further, the 3D structure-based distance measure 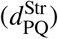 is also positively correlated with domain composition and duplication distances (Pearson *r* ≥ 0.5).

Next, to learn an emergent distance measure, *D*_PQ_, we combined four individual distances 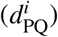, representative of E3 sequence, domain composition, structural, and functional level organization, with their appropriate weights (*w*_*i*_ ∈ {0.05, ···, 0.95} in 0.1 intervals). By uniformly sampling the weights, we constructed 10^5^ combination measures as a function of the hyper-parameter (fractional tree cutoff, *h*, between 0.05 and 0.95). By simultaneously maximizing element-centric similarity of the emergent hierarchical clusters resulting from combined measures, with partial ground truth (weakly-supervised scheme, **Fig. 2d**), we optimized an emergent distance measure (*D*_PQ_) with appropriate weights (ŵ _*i*_). We found that the linear combination of distances provided clusters with high element-centric similarity *S*_EC_ compared to clusters obtained from individual distances (**Fig. 2e**, black curve vs. colored).

Normalized Mutual Information (NMI) and Fowlkes–Mallows Index (FMI) compare clustering assignments (various distance-based vs. ground truth), but they are sensitive to cluster count (determined by tree cutoff, *h*; **Fig. S3a**). Therefore, optimized weights ŵ _*i*_ were obtained by averaging one hundred realizations of hierarchical clustering with maximum *S*_EC_ (*22*). The weights corresponding to maximum *S*_EC_ initially varied and then plateaued (at *h* ≥ 0.75; **Fig. 2f**), resulting in the construction of an optimized emergent distance measure, *D*_PQ_ (**Eq. 1**). We found that the relative influence of 3D structure, domain composition, and sequence alignment was more significant on the final learned metric and its ability to reproduce class labels accurately. Compared to the emergent distance measure, we found variable tree topologies with poor overlap and highly entangled trees for all four individual distances (**Fig. S3b–e**).

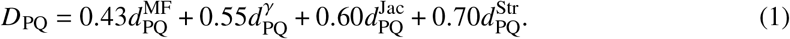

### Organization of the human E3 ligome

Using the optimized emergent distance metric, *D*_PQ_ (**Eq. 1**), we constructed a scaled hierarchical tree classifying the human E3 ligome (**Fig. 3** and **Fig. S4a**). To assess the validity of nodes, branch stability, and the robustness of our classification, we resampled the emergent distance matrix (*n* = 500) and assigned bootstrap support at each branch point (**Fig. 3**, grey circles). The bootstrap support for all nodes beyond tree cutoff, *h* > 0.15, is 95–100%, indicating a stable branch pattern (**Fig. S4b**) with a fixed tree topology. At *h* ≤ 0.15, the bootstrap support for the nodes dropped drastically. This allowed us to use a tree cutoff threshold, *h* = 0.25, to parse the dendrogram and obtain robust and stable clusters with clear family and subfamily patterns while preserving RING-, HECT-, and RBR-class segregation.

**Figure 3:**
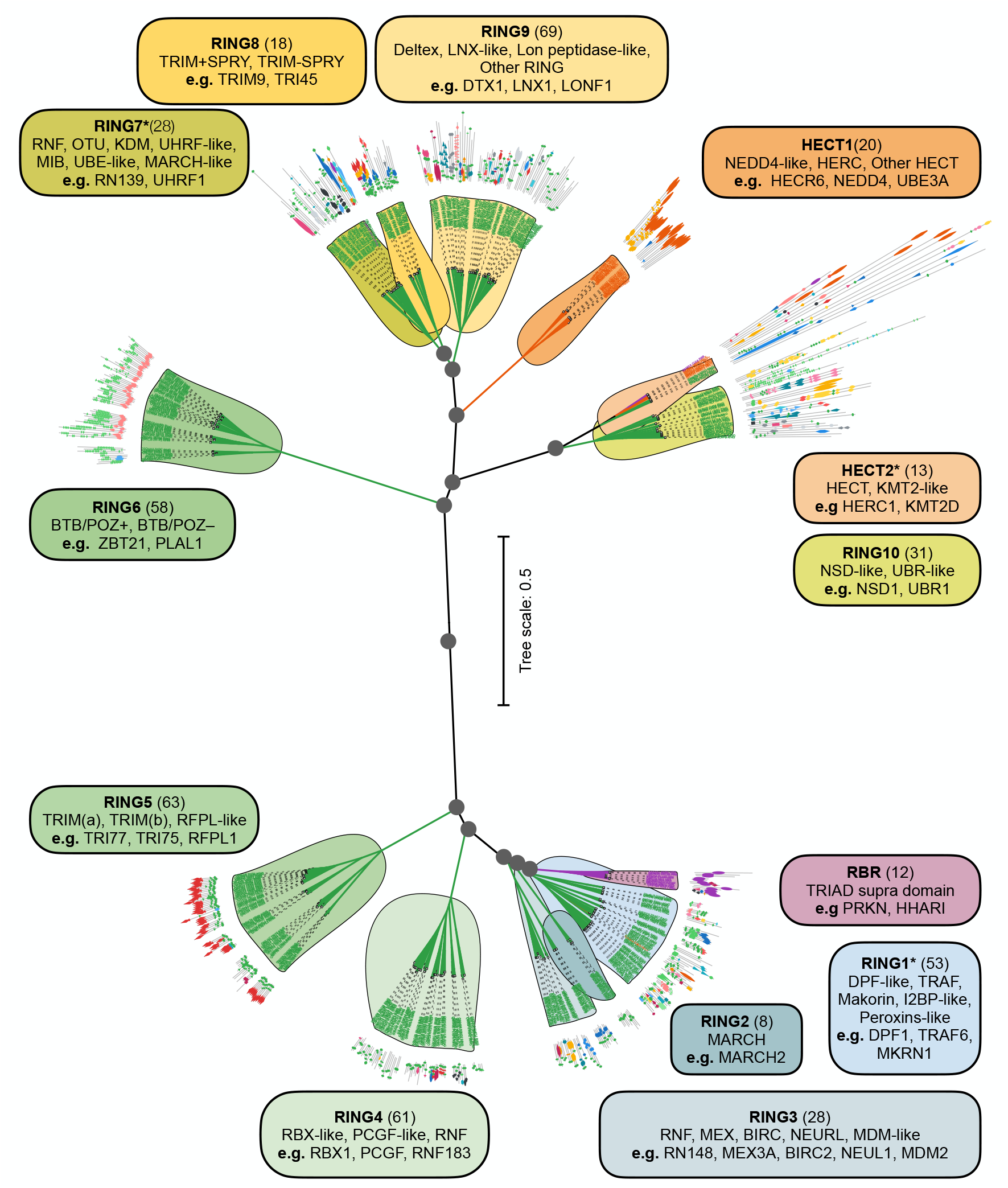
Classification of the human E3 ligome. Unrooted hierarchical tree computed using the optimized emergent distance metric *D*_PQ_ (scaled branch lengths). The RBR (purple), HECT (orange), and RING classes (blue/ green/ yellow) are partitioned at *h* = 0.25 into 1, 2, and 10 families, respectively. Each cluster is defined by shared sequence, domain-architectural (mapped), structural, and functional elements. Boxes show family information, i.e., family name, size, and subfamilies, with representative examples. Grey-filled circles denote bifurcation nodes with ≥ 95% bootstrap support, and * denotes families with a few class-level outliers (3/13).

We identified thirteen distinct clusters or E3 families (*h* = 0.25). At the class level, the E3 ligome is well segregated into ten RING families (**Fig. 3**, blue to green colors; clock-wise arrangement from RING1 to RING10), two HECT (**Fig. 3**, top-branch; orange), and one RBR family (**Fig. 3**, bottom-branch; purple). Each E3 family is subdivided into one or more subfamilies (**Fig. 3**, boxes) with distinct patterns. Mapping domain architecture information onto the individual leaves aids recognition of well-preserved sequence and domain features, consistent with family and subfamily grouping, a pattern more evident in the unscaled circular dendrogram of the E3 ligome (**Fig. S4a**).

Further, few heterogeneous families are grouped more closely and emerge from single branches (bootstrap support ≈ 90–95%; **Fig. S4b**) hinting at divergence of plausible superfamilies: (i) RBR and RING1–3 branch (small E3s), (ii) RING7–9 branch (medium E3s), and (iii) HECT2–RING10 branch (large E3s). This organization stems from the central node that bifurcates the E3 ligome into two groups characterized by average protein size (**Fig. 3**). The bottom branch displays six families with smaller E3s, while the top branch groups seven larger E3 families.

E3 family organization reflects mechanistic differences. The RING E3s mediate the direct transfer of Ub to the substrate, while the RBR and HECT E3s enable ubiquitin transfer via a twostep mechanism (**Fig. S4c**). The RBR-containing E3s form a homogeneous cluster, highlighting their conserved sequence and the TRIAD supra domain. Similarly, HECT domain-containing E3s are organized into two clusters/families, HECT1 and HECT2. The HECT1 family is homogeneous and includes three subfamilies: NEDD4-like, HERC, and other HECT E3s. The HECT2 family contains a pure HECT E3 subfamily and an outlier subfamily containing large multi-domain RING-type E3s that exceed 2000 amino acids in length. The most abundant RING-domain-containing E3s are organized into 10 families, each characterized by further grouping related proteins into distinct subfamilies with shared sequence elements, domain architectures, and structural features (**Table S3**). For instance, the RING2 family comprises membrane-associated RING-CH-type domain (MARCH) E3 ligases (**Fig. 3**, bottom-right). This family includes all small MARCH E3 ligases characterized by their transmembrane domains and sequence lengths below 500 amino acids. TRIM E3 ligases are exclusively limited to two distinct families, RING5 and RING8, and feature the SPRY domain (**Fig. 3**, bottom-left). E3 ligases containing BTB/POZ and Zn-finger domain repeats are grouped into the RING6 family (**Fig. 3**, upper-left).

Although our emergent metric largely maximizes pure and homogeneous clusters (e.g., RBR, RING2, RING5, RING6, RING8, and HECT1), heterogeneity often arises at the subfamily level, resulting in sub-groupings of E3s with varied and unique domain architectures. Isolated proteins (singletons) in the RING1, RING7, RING8, and RING9 families form distinct subfamily groupings, complicating pattern detection. Only RING1, RING7, and HECT2 families display occasional class-level outliers (**Table S3**). Supplementary Texts S1 to S13 describe each family structure in detail with information on subfamily branching, characteristic features, and distinct patterns along with outliers providing a nuanced description (**Figs. S5–S18** and **Supporting Texts S1–S13**).

### Functional segregation of the human E3 ligome

To understand the functional diversity of the human E3 ligome, we performed GO enrichment analysis and mapped our ligase classification and family structure onto it. This enabled us to draw clusters with unique functions and visualize their networks across all three ontologies. Further, mapping individual E3 ligases to these functions recognized the generic and family-specific functions.

At the biological process level, as expected (**Fig. 4a**), the network analysis revealed prominent core functional subclusters associated with all terms containing “ubiquitination (Ub)”, such as Ub-related processes, protein Ub, poly-Ub, K63-linked Ub, and positive regulation of catabolic processes (**Fig. 4a**, right bottom). These processes are shared across all families, indicating their essential roles in protein modification and degradation pathways. Another significant core functional cluster is centered around the innate immune response and regulation of type-I interferon production (**Fig. 4a**). In addition, the network highlights specialized functions like DNA metabolic processes and ERAD pathway regulation, demonstrating the diverse roles of E3 ligases beyond their canonical functions. The interconnectivity between GO functional clusters indicates cooperation across different biological processes by E3 systems. This is particularly evident for enriched functions involved in regulatory processes: regulation of type-I interferon production, regulation of response to biotic stimulus, regulation of defense response to virus, suppression of viral release by host, innate immune response, regulation of canonical NF-*κ*B signal transduction, and positive regulation of autophagy—all connected to protein modification and positive regulation of catabolic processes.

**Figure 4:**
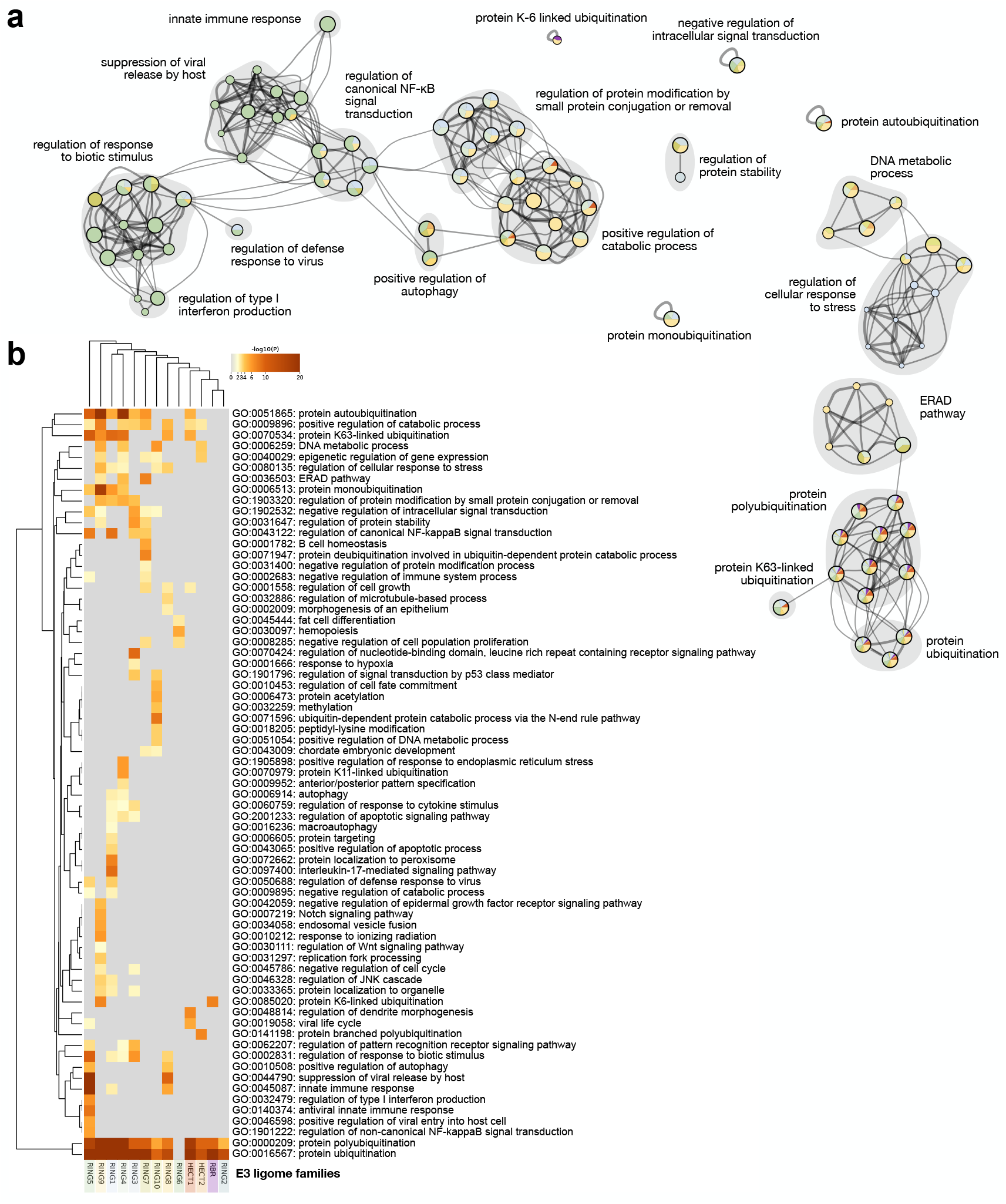
Functional segregation of the E3 ligome. **(a)** The functional landscape of the E3 ligome (biological processes) is captured by the network with GO annotation clusters. Individual nodes representing GO clusters (20 labeled) are drawn as pie charts (size *∝* number of E3s; colored by family enrichment) connected by distinct edges (*κ*-similarity ≥ 0.3). **(b)** The heatmap displays all functional clusters corresponding to family-specific enrichments of E3 ligases (discrete color scale for *p*-value *≤* 0.01; grey otherwise).

The analysis of E3 family-specific biological processes revealed distinct patterns of enrichment. For instance, RING5 E3s are enriched in regulating antiviral response, type-I interferon production, regulation of viral entry, and NF-*κ*B signaling. Similarly, RING8 E3s regulate innate immune response by suppressing viral release and positively regulating autophagy. RBR family E3s specialize in K6-linked ubiquitination, whereas the HECT2 E3s are responsible for branched polyubiquitination. We identified over 60 biological processes enriched with E3s corresponding to distinct families (**Fig. 4b**).

Distinct subcellular localization of E3 ligases directly exerts spatial control of ubiquitination (**Fig. S31a**). Most E3 ligases are cytosolic, which form an essential part of the ubiquitin ligase complexes (Generic function). Our analysis showed that the RING1 family members are enriched in the CD40 receptor complex, GID complex, and nBAF complexes; RING2 E3s are associated with early endosomes and lytic vacuoles; RING10 E3s are predominantly present in SWI/SNF complexes, associate with histone acetyltransferases and the nuclear chromosome; and RING9 members are associated with PML bodies, nuclear speckles, sites of DNA damage and ER quality control compartments. We identified 20 unique cellular components with distinct E3-specific enrichment patterns (**Fig. S19a**).

At the molecular level, all E3s are involved in ubiquitin-protein ligase activity (Generic function; **Fig. S19b**). This is often related to modification-dependent protein binding and ubiquitin-like protein binding, revealing key variations of enzymatic and binding activities catalyzed by E3s. The Zn-finger domains of RING E3s are responsible for engaging the E2–Ub complex and are also common to transcription factors. They could mediate chromatin binding, histone modifications, helicase activity, and unmethylated CpG binding functions. Alternate molecular functions of E3s stem from the extensive repertoire of domains and their unique family-specific domain architectures. They equip E3s to carry out diverse molecular functions such as p53 binding (RING3), ubiquitin conjugation (RBR), histone ubiquitination (RING9), unmethylated CpG binding (RING7), cullin family protein binding (RING4), etc. More than 25 molecular functions could be attributed to unique E3 family-specific domain organizations (**Fig. S19b**).

### Interaction landscape of the human E3 ligome

E3 ligases can operate as standalone or complex multi-subunit enzymes. In complex mode, E3 ligases are part of large multi-subunit complexes, including scaffold proteins, substrate receptors, and adaptors that support varying specificity, stability, and regulatory functions (*21*). For example, the Ring-box protein 1 (RBX1) is a core component of cullin-RING ubiquitin ligases (CRLs) essential for structural assembly and activity (**Fig. 5a**). RBX1 binds to the cullin scaffold proteins (CUL1–CUL5) and anchors the E2 enzyme, forming the crucial catalytic core of the complex to transfer ubiquitin to substrate proteins. The interaction of RBX1 with different cullins, substrate adaptors, and receptors allows for multiple CRL configurations (∼ 250), which provide modular regulatory control and confer specificity to diverse substrates.

**Figure 5:**
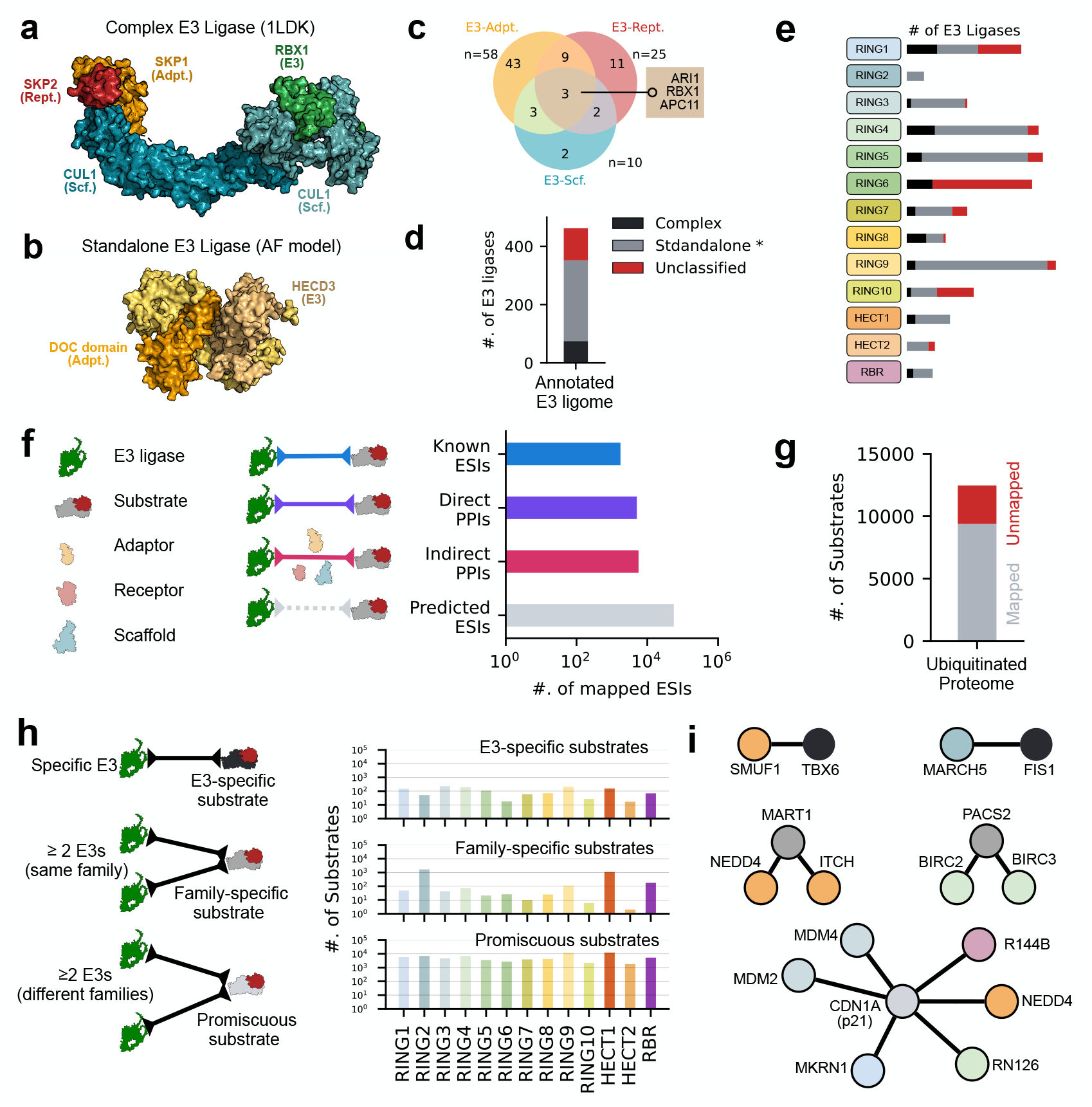
Protein–protein interactions of the E3 ligome. Representative examples of E3 ligases functioning as a **(a)** multi-subunit protein complex (CRL) or **(b)** a standalone enzyme (HECD3). Venn diagram of pairwise interactions of adaptors, receptors, and scaffold proteins with E3s. Annotation of 462 E3 ligases into complex, standalone, or unclassified modes of action. **(e)** Family-wise mapping of data from **d. (f)** Pairwise E3–substrate interactions for all E3 obtained by integrating data from known ESIs, mapped transient direct and indirect PPIs and predicted ESIs. **(g)** Mapping of the ubiquitinated proteome with E3s (≈ 75%, *n* = 12464). **(h)** Schematic showing substrate categorization into E3-specific, family-specific, and promiscuous classes (left) and their relative distributions mapped onto E3 families (right). **(i)** Representative examples for the three types of ESI networks.

By contrast, standalone E3 ligases, like MDM2, c-CBL, PARKIN, or SMURF1/2, either have specialized domains or undergo specific PTMs that recognize substrates and facilitate E2 binding and ubiquitin transfer. For example, HECTD3, like other HECT domain ligases, operates via a two-step ubiquitin transfer mechanism (**Fig. 5b**). However, substrate binding occurs through specific motifs within the non-HECT regions (DOC domain) that serve as adaptors and presumably recognize particular sequence motifs, distinct PTMs (e.g., phosphorylation), or unique structural elements of substrates.

Previous annotations (*23,24*) reported 6 E3s forming multi-subunit complexes (FBX30, KDM2A, FBX40, FXL19, KDM2B, and FBX11), 329 standalone E3s, and several unclassified. By integrating disparate interaction data, we extended this annotation. We first curated adaptors (*n* = 144; e.g., GAN, KLH21, SPOP), receptors (*n* = 91; e.g., SKP2, ASB3, CISH), and scaffold (*n* = 9; e.g., CUL1, ANC2, CACL1) proteins and cataloged their direct physical interactions with E3s (**Fig 5c**). The holo complex structure is only resolved for three E3 ligases (RBX1, ARI1, and APC11). There are 12 E3s with partial complex structures (APC11, ARI1, ARI2, KDM2A, KDM2B, PCGF1, PPIL2, PRP19, R113A, RBX1, RBX2, ZBT17). However, we found several binary direct physical interactions between E3-adaptor, E3-receptor, and E3-scaffold proteins, re-annotating 75 E3s operating in a complex mode (**Fig. 5d**, black), leaving 277 standalone E3s (*23*) and 110 unclassified E3s (**Fig. 5d**, red). Mapping this information onto the E3 ligome revealed that the RING8 family displayed the highest percentage of complex E3s (50%) followed by RING1 (26%), while RING2 and HECT2 families displayed entirely standalone E3s (**Fig. 5e, Table S4**). Consistent with our findings, we observe that MARCH-type E3s (RING2) operate in the membrane environment primarily as standalone enzymes. Further, the HECT2 family contains large multi-domain proteins with explicit domains to compensate for adaptor, receptor, and scaffolding functions (e.g., HECD3), explaining their standalone mode of action.

Next, we constructed the E3–substrate interaction (ESI) network by integrating data from known ESIs (*n* = 2012; known ESI; UbiNet + UbiBrowser), direct protein-protein Interactions (PPIs) (*n* = 5844; Direct PPI; IntAct DB), indirect PPIs (*n* = 6528; indirect PPIs; IntAct Db), and predicted ESIs (*n* = 64802; Pred. ESI; UbiBrowser pred., Top 1%). Integrating these data (**Fig. S19a**) by filtering high-confidence interactions (**Fig. S19b**) and verifying their ubiquitination status (overlap with PhosphoSitePlus or dbPTM) resulted in excluding false positives (E3-associated proteins) and improving the annotation of likely substrates (**Fig. S19b**). This enabled mapping ≈75% substrates (*n* = 9385 / 12464 proteins) from the ubiquitinated human proteome (**Fig. 5g**). Analysis of the E3–substrate network revealed distinct specificity patterns. Using well-known

ESIs alone, we found that the distribution of the number of substrates per E3 ligase is skewed. Several E3s have only one substrate (∼10^2^), some E3s target multiple substrates (∼10^1^), and very few E3s have an extensive portfolio of substrates (**Fig. S19d**). Given that a significant proportion of the proteome is ubiquitinated by the E3 ligome (462 E3s), most substrates are ubiquitinated by E3s belonging to two or more families (*n* = 7256 Promiscuous substrates; **Fig. 5h**; **Table S5**). However, we also identified substrates that are potentially ubiquitinated by two or more E3s belonging to the same E3 family (*n* = 3292 Family-specific substrates; **Fig. 5h**) and substrates uniquely targeted by specific E3 ligases (*n* = 1369 E3-specific substrates; **Fig. 5h**).

For instance, the E3 ligase SMUF1 specifically targets TBX6 for degradation during cell differentiation (*25*). Similarly, MARCH 5 specifically targets FIS1 for ubiquitination (**Fig. 5i**) to regulate mitochondrial fission (*26*). Both NEDD4 and ITCH belong to the HECT family and ubiquitinate MART1 to exert complementary functions for the sorting and degradation (*27*), and PACS2 is ubiquitinated by BIRC2 and BIRC3, members of the RING3 family (**Fig. 5i**), conferring TRAIL resistance to hepatobiliary cancer cell lines (*28*). CDN1A (p21), an essential factor in controlling cell cycle progression and DNA damage-induced inhibition of cellular proliferation, functions as a ubiquitous substrate. Several E3 ligases, such as MKRN1 (RING1), MDM2, MDM4 (RING3), RN126 (RING4), NEDD4 (HECT1), and R144B (RBR) families, target it, thus integrating several signaling pathways into replication checkpoints (**Fig. 5i**).

### Druggability map of the human E3 ligome

To learn likely avenues of proximity-based therapeutics and leverage the relationships within the human E3 ligome, we first mapped existing E3 handles derived from known Proteolysis Targeting Chimeras (PROTACs) and E3 binders to individual E3s and their families (**Fig. 20a, Table S6**). Only 16 proteins (9 catalytic E3s and 7 adaptors) are directly targeted by existing E3 handles (**Fig. 6a**, top). A large fraction of the designed E3 handles are specific to adaptor proteins (VHL, CRBN, DDBI, ELOC, KEAP1, DCA15, and KLH20), and a very select few directly target the catalytic E3s (BIRC2, XIAP, MDM2, BIRC3, BIRC7, RN114, UBR1, MDM4, and RNF4). We quantified the nearest neighbors for these nine E3s within RING3, RING4, and RING10 families and found an additional five closely related proteins (BIRC8, RN166, RN181, RN141, and UBR2; **Fig. 6a**, top; grey boxes). Given their high structural similarity (often paralogs), the same E3 handles could be repurposed to target them. Data on other family or protein-specific E3 handles are unavailable in the public domain. Mapping small-molecule E3 binders gave us a potential set of new lead compounds for the rational design of new E3 handles. We mapped E3 binders for 26 additional E3s and 15 auxiliary proteins (adaptors, receptors, and scaffold proteins), thus identifying new target proteins and avenues for lead development for the rational design of E3 handles (**Fig 6a** bottom; red labeled).

**Figure 6:**
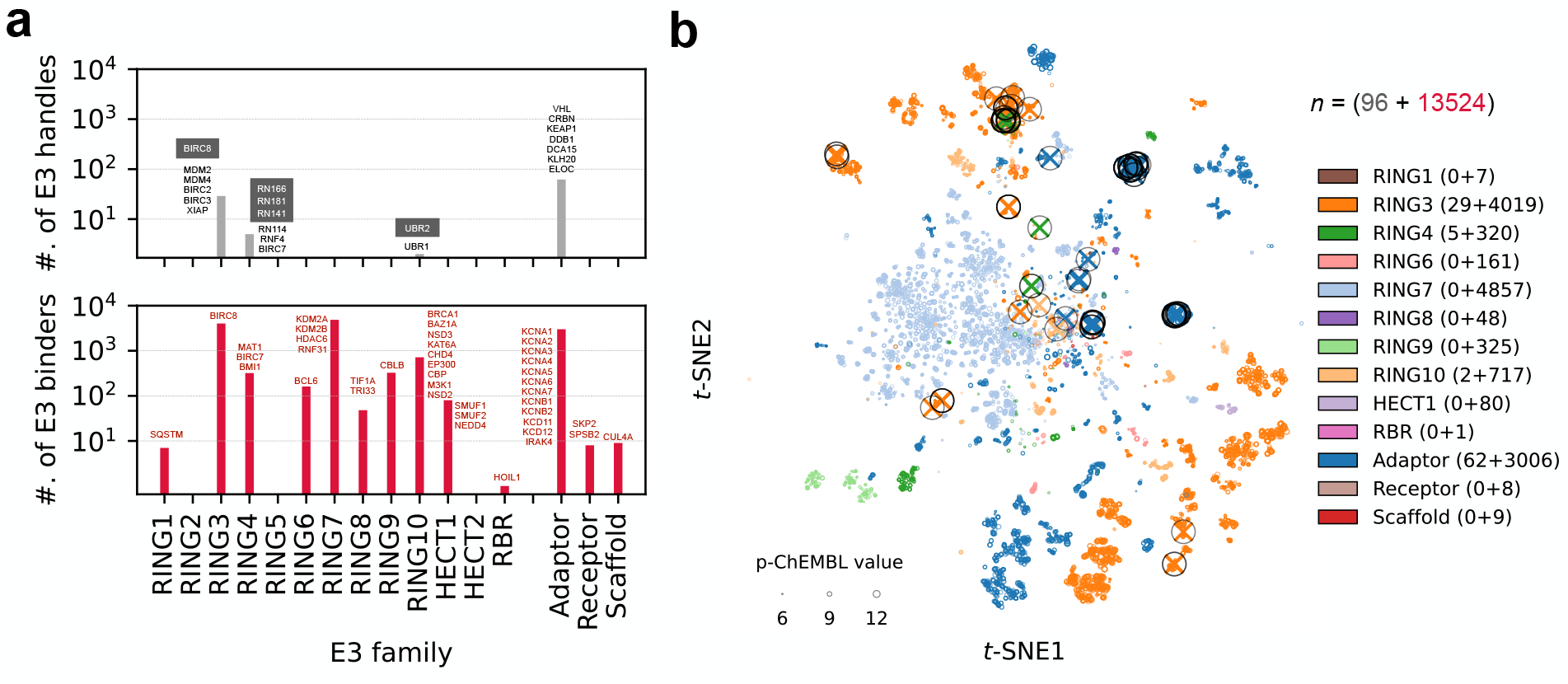
Druggability of the E3 ligome. **(a)** Distribution of known E3 handles (extracted from PROTACs, top) and newly identified E3 binders (potential lead compounds, bottom) targeting E3 families. Individual proteins uniquely targeted by E3 handles (*n* = 16, black) and E3 binders (*n* = 41, red) are displayed for each family. Grey-filled boxes (top) show closely related protein targets for E3 handle/PROTAC re-purposing. **(b)** Reduced chemical space using t-SNE showing the clustering of family-specific E3 handles (*⊗*) and unexplored E3 binders (O; Circle size *∝* p-ChEMBL value).

Next, we mapped the chemical landscape of E3 handles and binders. Using the *t*-distributed stochastic neighbor embedding (t-SNE) of high-dimensional 2048-bit Morgan fingerprints, we visualized their molecular similarities (**Fig. 6b**). We detected several chemically distinct clusters within the t-SNE subspace, targeting specific E3 families (distinct colors). E3 binders specific to RING3 (orange), RING7 (light blue), and adaptors (blue) occupy a large region of the chemical space, forming multiple dense clusters. Protein-wise decomposition of these E3 family-specific clusters revealed chemically distinct chemotypes within individual binder groups (**Figs. S20–S24**). For several clusters targeting RING3, RING4, and adaptor proteins, an E3 handle is often prominent and close to the representative E3 binder, indicating that the immediate chemical neighborhood represented by binders has characteristics specific to the given E3 (**Fig. S20–S21**). Further, the cluster density estimates the local sampling of chemical groups on central chemical scaffolds. (see examples for RING3, RING4, and RING10 families, **Figs. S20–25**). Furthermore, multiple protein-specific clusters within the t-SNE subspace indicate distinct pharmacophore fingerprints corresponding to alternate protein-small molecule binding sites. For instance, among adaptors, IRAK4 has six distinct chemical scaffolds, while KCNA5 and KEAP1 have 3 distinct scaffolds each (**Fig. S20c**). Similarly, MDM2 and XIAP (RING3 E3s) have five chemically distinct clusters specific to each protein often shared with closely related paralogs MDM4, BIRC3, and BIRC8 (**Fig. S21a**).

## Discussion

Navigating the vast and complex landscape of E3 ligase biology requires a comprehensive approach. Despite decades of dedicated investigation, the intricate diversity and functional complexity of E3 ubiquitin ligases continue to pose a significant challenge. In decoding this complexity, we first curated and filtered E3 ligases, ensuring data accuracy, consistency, and relevance for all downstream analyses. By assigning confidence scores to each ligase and employing stringent inclusion criteria, we remove false positives and improve annotation, providing a high-quality and comprehensive human E3 ligome. Ultimately, this simplification facilitated the identification of key catalytic components and paved the way for applying machine learning and algorithmic approaches to E3 systems.

The human E3 ligome exhibits remarkable heterogeneity, evident in its diverse sequence, domain architectures, structures, and functions. This diversity is shaped by not only the evolutionary forces influencing domain shu?ing and genetic rearrangements but also biophysical forces influencing molecular recognition and spatiotemporal regulation of enzymatic reactions, leading to specialization and adaptation (*29*). To effectively categorize E3 ligases, we require overarching organizational principles delineating broad evolutionary clans and functionally distinct subgroups within the E3 ligome. Hierarchical classification captures organizational principles, achieves higher prediction accuracy, and can handle novel data and class imbalances more effectively (*30*). These methods enable a more precise and context-aware organization of proteins, facilitating the recognition of salient and unique features (*31*). However, its performance heavily depends on choosing an appropriate metric reflecting authentic relationships.

Assessments of similarity and distance are critical components of human cognitive function and constitute a foundational element in developing and applying machine learning and data mining techniques (*32*). Using a weakly supervised learning paradigm, we optimized a linear metric that is simple, scalable, and straightforward to interpret with broad applicability. We bridged the molecular scale from protein sequence, domain architecture, 3D structure, and molecular function, resulting in a unique measure capable of detecting subtle shifts, reproducing class-level grouping of E3s, and improving family and subfamily definitions.

We present a multi-scale classification model to analyze the human E3 ligome comprehensively. We identified thirteen distinct E3 families. Shared domains, comparable architectures, and similar 3D structures often explain their clustering into families and subfamilies. Our classification method offers a novel approach, moving beyond traditional taxonomic methods and subjective, ad hoc classifications. Although not explicitly dependent on any individual distance measure, it is strongly associated with shared structural similarities and domain architectures, providing exceptional resolution into functional specialization and mechanistic action of E3s.

The RING E3 ligases form the largest class, are grouped into 10 families, and display a striking diversity. Our analysis uncovered family- and subfamily-specific features, contributing to their unique placement within the E3 ligome. RING2, RING5, and RING9 families show significant enrichment in specific cellular components such as lytic vacuoles, cytoplasmic stress granules, and DNA damage sites, respectively, mediating distinct biological processes. All TRIM E3 ligases are grouped into RING5 or RING8 depending on their domain architecture (*33*). These findings offer new frameworks for exploring the diversity of E3 ligase functions under multiple cellular and disease contexts. For example, TRIM E3 ligases are often involved in neuronal homeostasis (*34*) (RING5 or RING8), along with MARCH E3 ligases (*35*) (RING2 family). The RBR class demonstrates remarkable homogeneity, suggesting strong evolutionary conservation (*36*). The HECT class is split into two individual families (HECT1 and HECT2), consistent with the previous classification (*37*). These organizational insights lead to interesting new hypotheses, revealing new roles for existing E3s in health and disease.

Given the scarcity of experimental data on E3 ligase functions, GO terms serve as proxies for function. GO term enrichment analysis showed that the principal generic functions of E3s, i.e., BP: involvement in ubiquitination, protein modification, protein degradation, CC: localization to E3 ligase complex or cytosol, MF: catalyzing the transfer of Ub, are preserved among all E3 families. Our classification scheme captures additional family-specific specializations of E3 systems, providing significant insights into the diverse biochemical and functional mechanisms regulated by individual families. For instance, the RING5 family showed considerable enrichment in immune response regulation, while the RING9 family demonstrated specialized roles in cellular stress response. RING2 are enriched in membrane-bound organelles, indicating their specialized roles in protein quality control and trafficking pathways. Specialized molecular functions correlate directly with enriched domains, such as histone or chromatin binding of RING 10 E3s containing PHD-type Zn-finger and SET domains (*38, 39*), and kinase binding of RING1 subfamily with MATH/TRAF domain (*40*).

Mapping the protein interaction landscape of the whole E3 ligome is challenging. We integrate disparate datasets to build enzyme-substrate network maps for each ligase family. We found that RING1, RING3, RING8, and RBR members display higher numbers of E3s operating as multi-subunit complexes, while RING2 and HECT2 members are believed to operate in a standalone manner, directly recruiting substrates. Further, we could classify substrate molecules into E3-specific, family-specific, and promiscuous substrates. Identifying E3-specific and family-specific substrates provides foundational data for understanding the molecular principles of substrate recognition. Recognition of shared patterns in substrates can point to a better understanding of individual E3-specificity and group-specificity of E3 families. Further, our ESI network can be enriched by orthogonal data on subcellular localization of E3s and substrates and cell- and tissue-specific expression patterns to explain the context-dependent regulation of E3s and the prevalence of promiscuous substrates.

Targeted protein degradation via PROTACs is a promising therapeutic strategy to target previously undruggable proteome (*41*). Despite its potential, progress in targeting new E3s and the rational design of new E3 handles has been gradual. Most often, PROTACs and glue-like compounds exploit ligands against well-known adaptor proteins like CRBN- and VHL-dependent modalities to target CRLs for specific degradation of substrates. Only a few E3s have been directly targeted using PROTACs (*42, 43*). By leveraging the E3 ligome structure, we extend the map of E3 handles, increasing the likelihood of repurposing existing PROTACs to target closely related E3s in a familyspecific manner. Further, by mapping entirely new E3 binders and associating them with new E3s, we build a curated set of lead compounds with unique chemical signatures for further rational design of novel E3 handles. Furthermore, exploiting the novel relationships offered by the E3 ligome, in combination with enriched ESI networks, functional analysis, and a list of already targeted and newly identified E3 binders, allows an efficient drugging strategy for unexplored targets.

In conclusion, the multi-scale classification framework developed here provides a comprehensive global view of the human E3 ligome. Mapping disparate multimodal and multi-resolution data onto the ligome structure, such as functions, interactions, and druggability, provides a systems-level understanding, enabling high-throughput screening and profiling. The metric learning paradigm developed here is simple and transferable to other areas of data-driven biology. We anticipate that the data and insights presented here will stimulate further research into E3 systems and drive the development of innovative therapeutics.

## Materials and Methods

### Building the human E3 ligome

We collected eight individual human E3 ligase datasets (*A*_1_, ···, *A*_8_) including previously published reports (*17–19*) and public repositories: E3Net (*24*), UbiHub (*23*), UbiNet 2.0 (*44*), UniProt (retrieved on 2023-02-13 with search keyword “e3 ubiquitin-protein ligase”) (*45*), and BioGRID (retrieved on 2022-01-26) (*46*) compiled using multiple distinct criteria (**Table S1**). We merged all of them to form an initial dataset 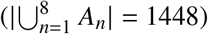, visualized the overlap of individual resources using UpSet plot (*47*), and assigned a consensus score to each entry based on its presence/absence among the source datasets. We then compiled a list of distinct, well-studied E3 catalytic domains from InterPro (*48*) corresponding to RING, HECT, and RBR classes from all published sources (*C* = {*d*_*C*_}; **Table S2**). Using the presence of characteristic catalytic domain(s) *d*_*i*_ within each polypeptide, we identified and filtered 1448 proteins corresponding to all catalytic subunits of E3 ligases,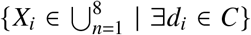. This was followed by manual curation based on InterPro domain descriptions of possible catalytic activity (E2-binding and Ub transfer) to obtain the final refined set of 462 E3 ligases (E3 ligome).

### Multi-scale distance measures

We encoded the pair-wise relationship of E3 ligases by computing twelve distinct distances (*d*_PQ_) spanning several granularity levels: primary sequence, domain architecture, tertiary structure, function, subcellular location, and cell line/tissue expression. All the distance measures were scaled between [0, 1] for comparison and even combination.

At the sequence level, we used an alignment-free local matching score-based (LMS) distance and an alignment-based γ distance between protein pairs using the canonical isoform sequences. The LMS distance 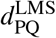 between two proteins P and Q is given by

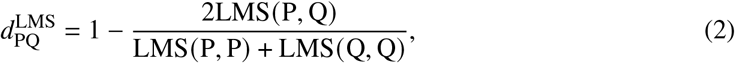

where LMS (P, Q) = ∑_*i*∈ {P,Q}_ *M* [*i, i*] captures the extent of local similarity by summing BLOS-SUM62 substitution scores for overlapping 5-residue fragment pairs {P, Q} from proteins P and Q (*49,50*). The pairwise γ distance measures the evolutionary distance between the globally aligned sequences of two proteins, P and Q, where *p*_PQ_ is the fraction of alignment positions with residue substitutions and indels, and *a* =2 (*51*).

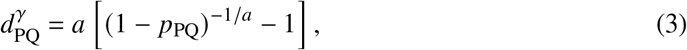

To quantify the preservation of domain architectures among all protein pairs, we computed three distances: Jaccard, Goodman–Kruskal γ, and domain duplication distances, using domain annotations obtained from InterPro database (*48*) (Nov 2022). The Jaccard distance (*52, 53*) represents the compositional similarity of protein domains. It is the ratio of the number of shared 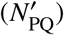 and unique domains 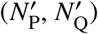 between proteins P and Q,

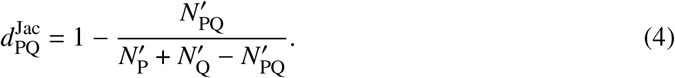

The Goodman–Kruskal γ distance compares the order of domain arrangements between two proteins, P and Q, and is computed as

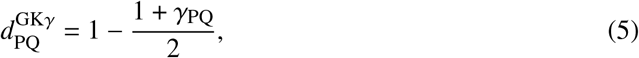

where 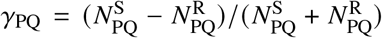 with 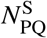 and 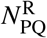 denoting the same- and reversed-ordered pairs of proteins P and Q, respectively (*53,54*). Finally, the domain duplication distance (*53*) compares the overlap of tandem domain repeats and is given by

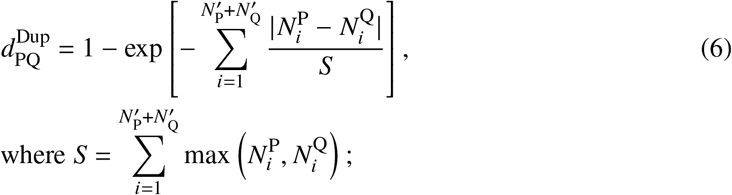

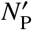 and 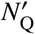 are unique domains in proteins P and Q with 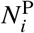 and 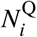 repeats, respectively.

To compute distances between structures of pairs of ligases, we used AlphaFold2 models (version 4) (*55*). We restricted comparisons to contiguous protein segments containing all catalytic domains for each protein to avoid comparing flexible regions of the full-length structures. We computed the TM-score as implemented in US-align (*56*). The TM score between the 3D structures of proteins P and Q is given by,

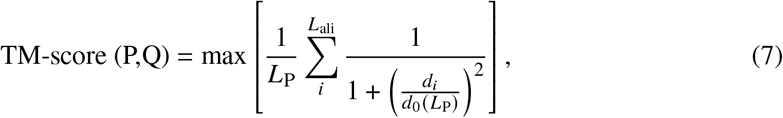

where *L*_P_ is the length of protein P, *L*_ali_ is the number of common residues between aligned proteins P and Q, and 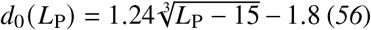 (*56*). To account for the inherent asymmetry in the TM similarity scores due to normalization by reference protein length *L*_P_, we computed the structural distance between protein structures P and Q by averaging their TM similarities as

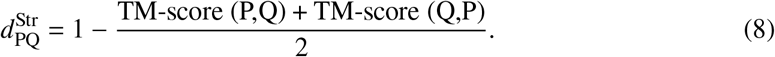

Functional distances among the protein pairs P and Q were captured using semantic similarities of annotated GO terms corresponding to the three GO ontologies—molecular functions, biological processes, and cellular components—using the package GOGO (*57*). The GO terms and the protein–GO-term mappings were retrieved (in Feb. 2023) from the Open Biological and Biomedical Ontology Foundry and the Gene Ontology resource (*31, 58*). For each annotated GO term x, we obtained a directed acyclic graph DAG_x_ = (x, T_x_, E_x_) with nodes T_x_ and edges E_x_. We defined the semantic contribution, following Wang et al. (*59*), *S*_x_(t) of a GO term t to the target term x as

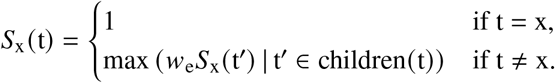

Further, the semantic similarity between two GO terms x and y, represented by two graphs DAG_x_ and DAG_y_, is defined as

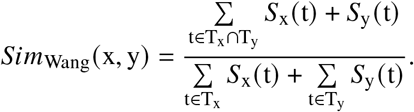

By extension, the semantic similarity between a single GO term x and a set of GO terms GO_Y_ = {y_1_, y_2_, ···, y_k_} is defined as the maximum semantic similarity between x and any of the terms in Y:

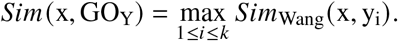

Finally, the semantic distance between proteins P and Q, annotated with sets of GO terms GO_P_ = {p_1_, p_2_, ···, p_m_} and GO_Q_ = {q_1_, q_2_, ···, q_n_}, respectively, is calculated as

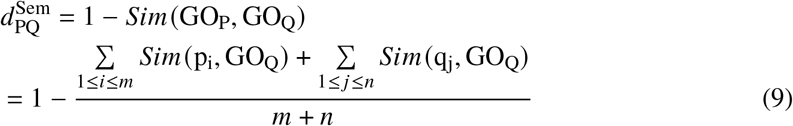

Using Eq. 9, we computed three semantic distances 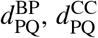 and 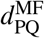 for the three different GO ontologies.

To compute the subcellular localization distance 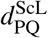, each protein’s main and auxiliary sub-cellular locations were mapped from the Human Protein Atlas (*60*) and used to construct a location vector with weights 1 and 0.3, respectively. We then computed 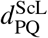 using the cosine similarity between the location vectors of proteins P and Q as

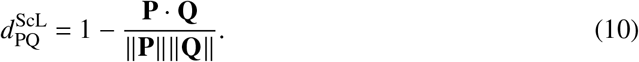

Finally, we computed the tissue 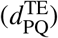 and cell line co-expression 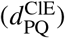 distances from the tissue and cell line expression profiles of the proteins P and Q. We retrieved expression data from the Human Protein Atlas (*60*), transcripts per millions of mRNA levels from the 253 human tissues of RNA HPA tissue gene dataset and 1055 cell lines of RNA HPA cell line gene dataset, respectively. Both distances were calculated using the Spearman’s rank correlation coefficient *r*_S,PQ_ as

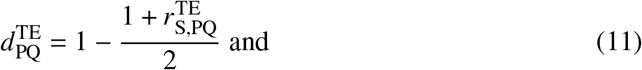

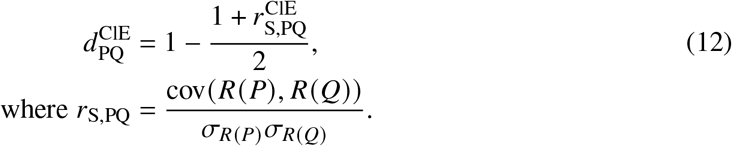

### Metric optimization, clustering, bootstrapping, and classification

We combined the pairwise gamma 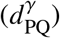, Jaccard 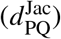, structural 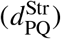, and semantic molecular function 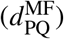 distances to capture all orthogonal information from the four significant hierarchies—sequence, domain architecture, 3D structure, and molecular function—into a single metric spanning the entire molecular scale. We used a weighted-sum model of these four distances, 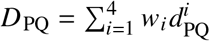, by uniformly sampling the weights as a function of tree cutoff, *h*, a hyperparameter. Optimized weights ŵ _*i*_, were obtained by maximizing the element-centric similarity index (*22*), which represents the similarity between clusters derived from parsing the emergent dendrogram (at evenly spaced cutoffs, *h* ∈ (0, 1)) derived from the combined distance and the class-level grouping of E3s into RING, HECT, and RBR classes (partial ground truth). At each cutoff *h*, we sampled ∼10^4^ emergent distance matrices (∑_*i*_ *w*_*i*_ *d*_*i*_), obtained their emergent hierarchical clusters, and computed *S*_EC_ for each one of them. We chose 100 emergent metrics with the highest *S*_EC_ for each *h* and computed the averages and standard deviations of their corresponding weights. The stabilized weights *ŵ*_*i*_ at *h* ≥ 0.9 corresponding to the maximum *S*_EC_ were chosen to construct the optimized distance measure. Dendrograms were computed from hierarchical clustering of individual and combined distance matrices using Ward’s minimum variance method (*61*) as implemented in SciPy. The emergent metric was resampled 500 times by swapping protein labels to compute bootstrap support at each bifurcation node. Unrooted trees with scaled distances were drawn and annotated with domain architectures of individual E3 leaves using iToL (*62*). The final tree was parsed at tree cutoff *h* = 0.25 to produce optimal emergent clusters (E3 families). Each family was manually analyzed for shared sequence and domain-architectural features to identify subfamilies and outliers.

### Identifying generic and specific functions of the E3 ligome

GO enrichment analysis for E3 ligases corresponding to individual 13 families was performed using Metascape (*63*), which implements a hierarchical clustering approach based on *κ*-similarity ≥ 0.3 (*63*). The resulting networks of GO terms at the biological process, cellular component, and molecular function ontologies were rendered using Cytoscape. Nodes were colored and drawn as pie charts to reflect E3 family contribution (number of proteins) and enrichment. Individual GO terms were considered significantly enriched within a ligase family if enrichment factor, *C*_obs._ / *C*_exp._ ≥ 2, a minimum of 3 proteins corresponding to the family are annotated explicitly with the corresponding GO terms, and a *p*-value ≤ 0.01). Within each resulting GO cluster, the GO term with the lowest *p*-value was selected as the cluster label for visualization. Heatmaps showing the enriched GO clusters for each family were drawn to highlight the functional specialization of individual E3 families.

### Integrating PPI and ESI datasets

To identify E3 ligases likely functioning in complex mode, we combined data from PDB (https://www.rcsb.org/) and IntAct (*64*). Using the refined lists of proteins corresponding to the E3 ligome (*n* = 462), E1, E2, adaptors, receptors, and scaffold proteins (Ubihub and manually curated lists), we retrieved all the PDB structures (as of Feb. 2023) involving E3-adaptors, E3-receptors, and E3-scaffold proteins. Following this, pairwise PPIs were obtained between E3-adaptor, E3-receptor, and E3-scaffold proteins filtered for “experimentally validated” PPIs (MI:0045) with high confidence (PSI-MI score ≥ 0.5). E3s interacting, or in a resolved structure, with at least one receptor, adaptor, or scaffold protein were re-annotated as complex E3s.

To construct E3–substrate interaction maps, we integrated multiple data sources, including experimentally validated enzyme-substrate interactions (ESIs) from UbiNet 2.0 (*44*) and UbiBrowser (*65*), a set of predicted ESIs from UbiBrowser (top 1% of predictions), physically interacting protein pairs (PPIs) from the IntAct database (mapped PPIs), and indirect PPIs involving ligases and potential substrates mediated by adaptor, receptor, or scaffold proteins from IntAct (indirect PPIs). Known ESIs and the PPIs dataset were enriched using substrates detected mainly by pull-down experiments, followed by two-hybrid techniques. A map of the ubiquitinated human proteome was obtained by cross-checking the ubiquitination status and mapping ubiquitination sites for each identified substrate from dbPTM (*66*) and PhosphoSitePlus (*67*). All substrates were categorized based on their interactions with E3 ligases: those paired with a single, unique E3 ligase were classified as E3-specific; those associated with multiple E3 ligases from the same family were designated as family-specific; and those linked to two or more E3 ligases from different families were labeled promiscuous.

### Mapping small molecule interaction data

A unified dataset E3 handles (corresponding to all publically documented PROTACs) and E3 binders targeting specific E3s, adaptors, receptors, and scaffold proteins were obtained by combining data from PROTACpedia (https://protacpedia.weizmann.ac.il), and PROTAC-DB 3.0 (*68*) and ChEMBL v34 (*69*). All small molecules were uniquely identified by their chemical structure represented using the canonical SMILES format and mapped to their target proteins and E3 families. Information from ChEMBL v34 was gathered using an SQL query combining compound data, experimental data, and target protein information and filtered using data from binding assays (p-ChEMBL value ≥ 6; equivalent to 1*μ*M binding).

2048-bit Morgan fingerprint (*70*) for each small molecule was obtained using RDKit (http://www.rdkit.org) (2048 bits array, radius= 3). Dimensionality reduction was performed using t-SNE using the Python Scikit-learn package (default parameters: perplexity=30, early exaggeration=12, n iter=1000, min grad norm= 10^7^, metric=euclidean, init=pca) and visualized by coloring all family-specific and protein-specific small molecule binders. The most representative compound for a given cluster targeting any specific E3 was identified as the compound with the highest average pairwise Tanimoto coefficient, computed using RDKit, with every other molecule in the same cluster.

## Acknowledgements

We thank Ivan Dikic, Stefan Knapp, Gerhard Hummer, Marcel Heinz, Varun Shah, and Matthew Shapira, along with all members of the PROXIDRUGs consortium, for their support and constructive discussion. We thank David Krause for system administration and the Center for Supercomputing, Goethe University Frankfurt, for computing time on the Goethe-HLR cluster.

## Author Contributions

Conceptualization: R.M.B.

Methodology: A.D., A.C., S.V.N, and R.M.B

Investigation: A.D., A.C., S.V.N., and J.E.

Data analysis: A.D., A.C., S.V.N., J.E., and R.M.B.

Visualization: A.D., A.C., S.V.N., and R.M.B.

Supervision: R.M.B.

Funding acquisition: R.M.B.

Writing—original draft: A.D., A.C., S.V.N, and R.M.B.

Writing—review & editing: A.D., A.C., S.V.N, and R.M.B.

## Competing interest

### Professional affiliation R.M.B

Head scientist (Computational Biomedicine), Frankfurt Competence Center for Emerging Therapeutics (FCET), Goethe Center for (high) technology (Go4Tec), Goethe University, Frankfurt am Main, Germany.

This manuscript reflects the views of the authors, and neither IMI nor the European Union, EFPIA, or any associated partners are liable for any use that may be made of the information contained herein.

### Data and materials availability

All data supporting the findings are provided in the Supplementary materials and additional data files.

